# The syntax-meter interface in spoken language and music: Same, different, or individually variable?

**DOI:** 10.1101/2023.03.08.531723

**Authors:** Eleanor E. Harding, Daniela Sammler, Sonja A. Kotz

**Affiliations:** Department of Neuropsychology, Max Planck Institute for Human Cognitive and Brain Sciences, Leipzig, Germany; Department of Otorhinolaryngology/Head and Neck Surgery, University Medical Center Groningen, University of Groningen, Groningen, The Netherlands; Research Group ‘Neurocognition of Music and Language’, Max Planck Institute for Empirical Aesthetics, Frankfurt am Main, Germany; Faculty of Psychology and Neuroscience, Department of Neuropsychology and Psychopharmacology, Maastricht University, Maastricht, the Netherlands

## Abstract

Considerable debate surrounds syntactic processing similarities in language and music. Yet few studies have investigated how syntax interacts with meter considering that metrical regularity varies across domains. Furthermore, there are reports on individual differences in syntactic and metrical structure processing in music and language. Thus, a direct comparison of individual variation in syntax and meter processing across domains is warranted. In a behavioral (Experiment 1) and EEG study (Experiment 2), participants engaged in syntactic processing tasks with sentence- and melody stimuli that were more or less metrically regular, and followed a preferred or non-preferred (but correct) syntactic structure. We further employed a range of cognitive diagnostic tests, parametrically indexed verbal- and musical abilities using a principal component analysis, and correlated cognitive factors with the behavioral and ERP results (Experiment 3). Based on previous results in the language domain, we expected that a regular meter would facilitate the syntactic integration of non-preferred syntax. While syntactic discrimination was better in regular than irregular meter conditions in both domains (Experiment 1), a P600 effect indicated different integration costs during the processing of syntactic complexities in the two domains (Experiment 2). Metrical regularity altered the P600 response to preferred syntax in language while it modulated non-preferred syntax processing in music. Moreover, experimental results yielded within-domain individual differences, and identified continuous metrics of musical ability more beneficial than grouping musicians or non-musicians (Experiment 3). These combined results suggest that the meter-syntax interface differs uniquely in how it forms syntactic preferences in language and music.

## Introduction

For many decades, research in language and music has compared how their structural properties are perceived. The milestone Shared Syntactic Integration Resource Hypothesis (SSIRH, Patel, 2003; cf. Slevc & Okada, 2015) proposed that the online integration of syntactic constituents into their preceding context is monitored by the same neural resources across domains. This approach has recently been criticized and is subject to current debate (Asano & Boeckx, 2015; Fedorenko & Shain, 2021; Slevc & Okada, 2015).

The domains also share that humans perceive hierarchical temporal relationships among stressed syllables and beats — described as meter in both spoken language (e.g., Prince, 1983) and music (e.g., London, 2012). Intuitively, while respective meters are on a gradient with some overlap, spoken language tends to be perceived as less temporally regular than music (Kotz et al., 2018; Patel, 2008). Syntactic and meter alignment also differs (see Hilton & Goldwater, 2021) in language and music, but becomes relevant for cross-domain syntax comparison. Necessarily, as different speakers may produce different meter for the same sentences (Dellwo & Wagner, 2003), the syntax does not change if spoken meter is altered^1^ (cf. Selkirk, 2011). Conversely, music requires specific metrical construction to preserve syntactic well-formedness rules (e.g., Lerdahl & Jackendoff, 1983). Strong metrical boundaries serve as harmonic anchors, and thus are aligned with where a sense of harmonic resolution is most prominent (Bigand et al., 1999; Boltz, 1993).

Considering the cross-domain differences in metrical regularity and how meter aligns with syntax, a question that emerges is, how do differences in meter influence syntactic processing? This question is especially important if there is a shared resource, integrating the syntactic elements as described by the SSIRH. A growing literature shows that temporal regularity in the signal — which is linked to metrical structure — can influence syntactic processing in language. Namely, increasing temporal regularity in spoken language benefits syntactic processing (Roncaglia-Denissen et al., 2013; Schmidt-Kassow & Kotz, 2008). This also appears to be the case in music: the tonality of temporally expected musical elements is judged more accurately (Bigand et al., 1999). However, to our knowledge, no study has directly addressed how metrical structure influences musical syntax processing or whether the influence of meter on syntax processing is comparable across both domains.

A natural extension of this question is whether everyone benefits similarly from metrical regularity in language and music with respect to syntax processing. Syntactic processing is subject to differences in working memory (WM) capacity at least in language (e.g., Bornkessel et al., 2004), and musicians are reported to have higher working memory capacity in general (Kraus & Chandrasekaran, 2010). This perhaps explains why WM differences are seldom reported in music syntax studies, where musician participants may be clustered at high WM capacity (see Yurgil et al., 2020 for a review on musical training and working memory). Musical ability is also reported to influence cross-domain syntactic processing (Harding, Sammler, & Kotz, 2019; Menon, 2016) as well as the detection of metrical properties in language (Magne et al., 2007; Marie et al., 2011) and music (Geiser et al., 2010). Syntax and meter processing are also influenced by individual differences in temporal perception, because temporal perception may relate to better neural tracking of stimuli that evolve over time (Harding, Sammler, Henry, et al., 2019; Skoe & Kraus, 2010) — with benefits for metrical and syntactic processing (Ding et al., 2015). Thus, we assessed individual differences in working memory, musical ability, and temporal perception as predictors of syntax perception in varying metrical contexts.

The current set of studies aimed to show whether syntax processing is similarly affected by metrical regularity in music and language. Moreover, individual differences in working memory, time perception, and musical abilities might predict the strength of this syntax-meter interaction. We conducted three experiments with syntactically complex, metrically regular and irregular melodies and sentences. We showed that both behavioral (Exp. 1) and EEG measures (Exp. 2) converge on the influence of metrical regularity on syntactic processes across both domains though likely for different underlying reasons, while time- and pitch-related auditory ability impacted syntactic processing in music only (Exp 3).

## Experiment 1: Behavioral study

### Syntax

Language and music^2^ are both structurally governed by a ‘syntax,’ or a set of guiding structural principles that native listeners implicitly understand (Chomsky, 1957; Ellis, 2008; Lerdahl & Jackendoff, 1983; Rohrmeier & Rebuschat, 2012). Considering similarities in the construction of syntactic dependencies in music and language (Gibson, 1998; Lerdahl, 2005) as well as neuroimaging data suggesting an overlap in how the brain responds to syntax across the domains (e.g., Maess et al., 2001; Patel et al., 1998; Sammler et al., 2013), the SSIRH (Patel, 2003, 2008) proposed that the neural resources underlying the online integration of syntactic elements into their preceding context are shared across music and language.

Recent challenges to the SSIRH (Perruchet & Poulin-Charronnat, 2013; Slevc & Okada, 2015) have observed that most empirical, cross-domain syntax studies use paradigms that compare well-formed syntax with syntactic violations, thus error detection mechanisms may be compared as opposed to syntactic processes per se. Therefore, the syntactic constructions comprising the current studies are relative clauses in language and relative keys in music, which create complex structures that tax syntactic integration processes rather than overt error detection processes.

In language, a relative clause modifies some element in a sentence and usually begins with a relative pronoun such as *who* or *which*. When a relative clause modifies a complex noun- or verb phrase, the relative pronoun ambiguously refers to elements in the complex phrase it modifies (e.g., typical garden-path sentences; Sanz et al., 2015). In languages such as German, verb conjugation can resolve relative pronouns that are ambiguous in number (e.g., *who* can refer to *they* (plural) or *she* (singular)), but are correct no matter which way the ambiguity is resolved (Example 1). Thus, concerning ambiguous relative clauses, the listener must build a complex syntactic structure that holds two different possibilities open until the ambiguity is resolved. The syntactic structure among words is guided by information such as gender or number, which can be contained in morphemes across word categories.

Example 1.

Da hinten arbeiten die Diener der Ärztin, die Ghent vor kurzem…

*Back there are working the servants of the doctor, who recently…*

…besuchten und mochten.

*…visited (plural) and liked (plural) Ghent*.

…besuchte und mochte.

*…visited (singular) and liked (singular) Ghent*.

In Western tonal music, a tonal key consists of 7 pitches that are organized around a central tone or ‘tonic’. In the key of C major, for instance, C is the tonic and the other tones are described in relation to the C. Importantly, successive pitches in each key are separated either by one or by two semitones. In major keys, the one-semitone intervals are between the third and the fourth (E and F in Figure 1) and the seventh and the eighth tone (B and C in Figure 1).

**Figure 1.**
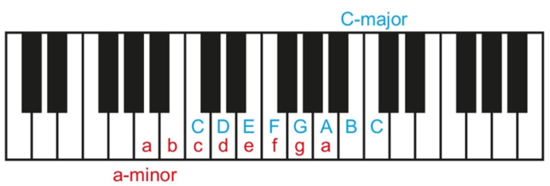
Relative tonal keys. The same set of notes comprise relative major and minor keys, for example C-major and a-minor shown here. The tonic differs (C or a, respectively) as well as the position of one-semitone intervals (between the 3rd and 4th, 7th and 8th tone in major; between the 2nd and 3rd, 5th and 6th tone in minor).

Each tonal key has a relative tonal key, which consists of the same set of notes but uses a different tonic as a reference point (a in Figure 1). With a different reference, the intervals between successive tones also change. For example, the one-semitone intervals now appear between the second and third- and fifth and sixth positions relative to the tonic. This characterizes the key as ‘natural minor’.

Thus, in a musical passage it is possible to shift between a relative major or natural minor key by changing the perceived tonic, without changing the tones themselves. This can be accomplished by, for example, placing the tonic of the relative key in prominent metrical positions, or creating melodic contours to highlight prominent major or minor intervals between consecutive notes. When a key changes within a passage, this is referred to as ‘modulation’ to a different key. In practice, the perceived tonal key in music is shaped by the intervals in the melodic contour and accompaniment, especially by notes placed at strong metrical locations, i.e., locations of structural salience such as the beginnings or ends of phrases or passages. Musical tension is created by moving the tonal structure further from the tonic. Syntactic well-formedness in music is related to how well tension in the passage is resolved, for example returning to the tonic at the end of a piece is the most common way to ‘relax’ or resolve any harmonic tension that may have been created in the interim (Lerdahl & Jackendoff, 1983; Piston, 1987).

### Meter

The perception of meter is inherent to both language and music. In language, within the study of phonology, one finds metrical theory and prosodic phonology that formalize linguistic aspects of spontaneous productions of native speakers. Here, we focus on the metrical grid theory (see Nespor & Vogel, 2012 for hierarchical organization in prosody; Prince, 1983). Metrical grid theory is concerned with the ultimate strong-weak relationships among speech syllables that extend across utterances (Prince, 1983)^3^. The emphasis on weak or strong syllables is adjusted by a speaker in order to preserve well-formed structure in the grid. For example, the word ‘Tennessee’ is pronounced with emphasis on the final syllable, (TennesSEE), and yet when pronouncing the words ‘Tennessee air’, native speakers shift emphasis to the first syllable (TENNessee AIR) in order to create a well-formed metrical grid with a strong-weak-weak-strong structure.

In music, meter is the abstract, hierarchical organization of beats that a listener perceives as music unfolds in time (London, 2012). Meter in music emerges with a combination of melodic contour of notes, relative length of notes and loudness changes (Hannon et al., 2004). The most typical Western meters have an alternating strong-weak beat perceptually emergent pattern, such that strong beats are grouped with two or three subsequent weak beats before a strong beat is repeated (e.g., strong-weak-weak; C. M. Wright & Simms, 2006). These smaller groups are organized such that the perception of larger groups emerges, which form metrical levels in an ascending hierarchy across passages and entire works.

The perception of language and music meter can be seen as occurring along a gradient from irregular to regular — regularity defined as a consistent ratio of strong and weak beats — such that the bulk of language meter construction typically occurs in less regular form than the bulk of music meter construction (Kotz et al., 2018; Patel, 2008). However, the variation in the two domains contains overlap, depending on the intention of the performer or speaker: for example, virtuosic improvisation or ornaments in musical passages might contain irregular timing, while a poetry slam or spoken religious liturgy might contain highly regular timing.

### Syntax and meter

An interesting point for cross-domain syntax processing that we note is the tendency for language meter to be less regular, and for musical meter to be more regular (Kotz et al., 2018). Moreover, meter can be flexibly arranged in language without necessarily altering the syntactic structure (Hilton & Goldwater, 2021), whereas in music meter and syntax are typically described as interwoven (Lerdahl & Jackendoff, 1983). Considering these differences in how meter and syntax align across the domains, one might surmise that syntactic resources across domains need to be different to account for the differences in meter.

Yet, despite these differences, single-domain paradigms have described similar effects of meter on syntax perception: regular meter facilitates the syntactic processing in some way. In language, syntactic integration is easier (shown by a reduced syntactic ERP effect) when syllables have a consistent strong-weak ratio in the meter as opposed to a meter that changes the strong-weak recurrence within a sentence (Roncaglia-Denissen et al., 2013) or as opposed to a meter that violates the metrical grid (Schmidt-Kassow & Kotz, 2009). In music, judging whether chords follow syntactic well-formedness is more accurate when the meter is regular as opposed to when strong and weak beat patterns are disrupted (Bigand et al., 1999; Schmuckler & Boltz, 1994).

To address how meter might affect language and music syntax processing, we created syntactically complex sentences and melodies with more- or less regular intended metrical composition. The syntactic complexity aimed to tax processing without triggering error detection mechanisms: determining relative clause attachment in sentences or modulation to a relative key in melodies. Regular meter repeated strong-weak-weak patterns and irregular meter shifted strong-weak patterns throughout. Based on previous single-domain studies (Bigand et al., 1999; Roncaglia-Denissen et al., 2013; Schmidt-Kassow & Kotz, 2008; Schmuckler & Boltz, 1994), we hypothesized that regular meter would facilitate syntactic processing in both domains.

## Materials and methods

### Participants

Thirty native German speakers (15 females, 20–31 years, *M* = 23.7 years, *SD* = 3.44) participated in this experiment. Participants reported a range of formal musical training from zero (non-musicians) to 18 years (*M* = 8.1 years, *SD* = 7.4) beyond obligatory school courses. A year of formal musical training was defined as a year during which the participant underwent a structured lesson schema of any instrument or voice, either self-taught or by an instructor, and practiced an average of at least one hour per week. All participants gave written informed consent prior to the experiment and were paid for their participation. The study was approved by the ethics committee of the University of Leipzig and carried out following the guidelines of the declaration of Helsinki.

### Stimuli

240 spoken sentences and 240 piano melodies with analogous construction were used in this study. Example stimuli are shown in Figure 2.

**Figure 2.**
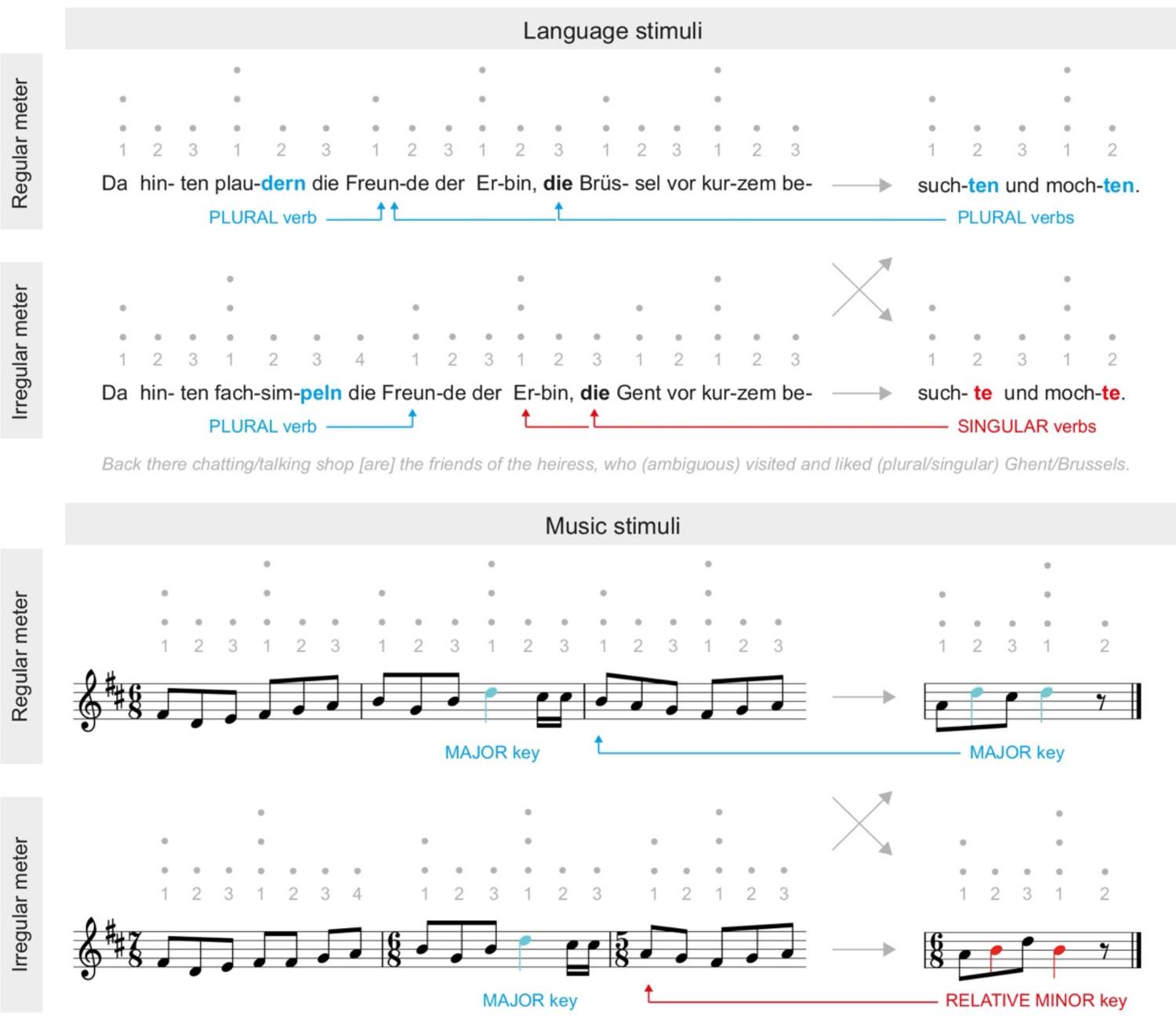
Language and music stimuli. *Upper panel:* Sentence stimuli consisted of a main clause (marked in blue) that established a plural subject (arrow to ‘die Freunde’) and a relative clause (relative pronoun ‘die’, in bold) that attached either to the plural subject or the singular genitive possessor (‘der Erbin’). This ambiguity was resolved in the plural or singular conjugation of the final verbs (cues marked in blue and red). Regular meter consisted of a strong syllable every three syllables, irregular meter shifted the strong syllable to create mixed groups of 2, 3, or 4 (see gray numbers). *Lower panel:* Melodies consisted of a first phrase in a major key (marked in blue) that was either confirmed or modulated to the relative minor key at the end of the second phrase (cues marked in blue and red). The metrical grid was analogous to the language stimuli. Inherently different syntax-meter alignments in music and language, however, required placement of the second critical cue in a metrically strong position in music while it occurred at a metrically weak position in language. Dots denote the metrical strength of syllables and notes according to grid theory.

### Language

Each sentence consisted of two clauses (see upper panel of Figure 2). In the first clause, plural verb conjugation clearly established a plural sentence subject (‘die Freunde’/*’the friends’*) that had a singular feminine genitive possessor (‘der Erbin’/*’of the heiress’*). The second clause started with an ambiguous relative pronoun ‘die’/*‘who’* that could attach to the subject (high) or the possessor (low) in the first clause. Disambiguation was first possible at the final verb conjugation (either ‘-ten’, plural subject or ‘-te’, singular possessor) at syllables 20 and 23 (see blue and red arrows).

The lexical content of the sentences was strictly controlled. Two prepositions started each sentence (ten combinations, e.g., ‘Da hinten’/*’Back there’*), followed by a verb conjugated for the first plural noun (five verbs total, paired with homonyms to create two or three syllables to fit the respective Meter condition, e.g., ‘plaudern’/*’are chatting’* and ‘fachsimpeln’/*’are talking shop’*), then an article with plural noun (15 animate words of relation: aunts, friends, etc.) to set up the first possibility for the ambiguous relative pronoun, next a genitive article and singular noun (15 animate words of a nationality, title or profession: heiress, customer, Englishwoman etc.), providing the second option for the ambiguous relative pronoun. In the second clause, the first item was always ‘die’/*’who’*, followed by a city (15 locations, either well-known international cities or cities within Germany, e.g., ‘Brüssel’/*’Brussels’,* which were paired with a similar class of city to create a one or two syllable variation: e.g., London and Leeds), followed next by the verb ‘besuchen’/*’to visit’* (conjugated for either singular or plural), followed by the conjunction ‘und’/*’and’* and a final verb, ‘mögen’/*’to like’* (conjugated to the same number as *‘to visit’*). Lexical items were pseudo-randomized to create 60 sentences with regular meter. Then a second set was reproduced with the partner three-syllable verb and one-syllable city in the first and second clauses to obtain 60 analogous sentences but with irregular meter. Finally, each of these 120 sentences was prepared once with plural and once with singular final verb conjugation, resulting in a total of 240 sentences. A native German speaker reviewed the sentences for grammaticality and plausibility. The lexical items for the sentences were controlled for frequency using the Leipziger Wortschatz and Celex word databases.

### Music

Each melody was composed of two phrases, each with two measures (see lower panel of Figure 2. The first phrase clearly outlined a major key (e.g., D-major, see notes marked in blue). In the second phrase, the tonal key either remained major (i.e., D-major) or resolved to the relative minor key (i.e., b-minor, see notes marked in red). Note that the major and its relative minor key contain physically identical notes making the melodies locally ambiguous (see blue and red arrows in Figure 2). Whether the key remained major or modulated to the relative minor key was first detectable at beat 20, which was consistently the tonic of this key (i.e., d in D-major and b in b-minor; see notes marked in blue and red, respectively), followed by repetition of the tonic at beat 22 via a final cadence in accordance with music syntax well-formedness rules (Lerdahl & Jackendoff, 1983).

Each of 12 melodies were presented in five different keys: C-major and its relative a-minor, G-major/e-minor, F-major/d-minor, Eb-major/c-minor, and D-major/b-minor, resulting in 60 major-major and 60 major-minor melodies. Each of these 120 melodies was prepared in a regular and irregular meter version, resulting in 240 stimuli in total. The keys were verified by a probabilistic key-finding algorithm (Temperley & Marvin, 2008).

### Alignment of syntax and meter

As mentioned in the Introduction, syntax-meter alignment differs between language and music. Grid theory in language (Prince, 1983) does not pose strict limitations on syntactic structure in terms of where relative pronouns or disambiguating verbs must occur. In fact these elements, though structurally important, often occur at weak metrical locations in German (see Figure 2). The tonal key in music, on the other hand, is more definitively shaped by notes placed at strong metrical locations, i.e., locations of structural salience such as the beginnings or ends of phrases or passages (Lerdahl & Jackendoff, 1983; Piston, 1987). Considering these different constraints, locations of the critical syntactic cues differed in their metrical position across domains. While the first cue was in a metrically weak position in both music and language (beat 20), the second cue was necessarily in a metrically strong position in music (beat 22) compared to a repeated weak position in language (beat 23). Hence, while it was always possible to use the first cue to determine the whole syntactic construction, the second cue might be more relevant in music.

### Fillers

T0 prevent participants from developing response strategies, additional filler sentences (*N* = 240) and filler melodies (*N* = 240) were presented in the experiment but excluded from further analysis. Fillers followed similar syntactic construction as the experimental stimuli and required integration of a syntactic cue with its context. However, instead of starting with an ambiguous pronoun, relative clauses in filler sentences began with a subject or object relative pronoun. Filler melodies, instead of modulating (or not) to the relative minor key near the end, modulated (or not) to another major key (e.g., a major second) at the beginning of the second phrase.

### Recording

The melodies and sentences were produced and recorded by a conservatory-trained pianist (30 years, female, 13 years of classical piano training) and a professionally trained native German speaker (30 years, female), respectively. Musical scores and sentences were given to the pianist and speaker several days before the recording sessions, to allow them to become familiar with the material. During recording sessions, a 100 beats per minute (bpm) metronome was provided to the pianist and speaker via headphones with the instruction to time the downbeats or strong syllables to the metronome. For the irregular meter melodies, the pianist requested a 300 bpm metronome and subsequently timed the subdivided beat to the metronome. The resulting syllable and note presentation corresponded to a relevant perceptual range in speech and music perception (Ding et al., 2017; Doelling & Poeppel, 2015; Luo & Poeppel, 2007). Melodies were recorded on a Yamaha Clavinova CLP 150 electric MIDI keyboard (Yamaha Corporation, Hamamatsu, Japan) with a 44.1 kHz sampling rate using Finale 2008 (Boulder, CO, USA). Sentences were recorded with a Rode NT55 microphone (Silverwater, Australia) at 16-bit resolution and a sampling rate of 44.1 kHz using Cool Edit Pro 2.0 (Sibiu, Romania). Recorded stimuli ranged between 4 and 5 s per item (mean 4.581 s, ± 0.0068 *SEM*). Peak amplitude (intensity) was normalized to 70 dB using Audacity 1.3 and Praat 5.2 (Boersma & Weenink, 2011).

### Task and procedure

The experiment was divided into a language session and a music session (see Figure 3) that were held on different days in counterbalanced order among participants. In each trial, participants were presented with two items from the same meter condition (regular vs. irregular) with 400 ms inter-stimulus interval (ISI). They were asked to compare the second item to the first and to indicate via button press as quickly and accurately as possible (i.e., speeded response task) whether the items were structurally the ‘same’ or ‘different’. This discrimination task required participants to detect the critical cues, which determined the larger syntactic dependencies in each domain. Trials were designed to be 50% same/different ratio and were presented in pseudorandom order such that same/different trials occurred not more than three consecutive times in a row. Regular and irregular meter trials did not occur more than three times in a row. ISI between trials was 1000 ms. The order of sentence or melody type (plural vs. singular verb, major vs. minor key) was counterbalanced across trials. Button assignment (‘same’/‘different’) on a computer keyboard was counterbalanced among participants for right and left hands. Participants completed a short training to make sure they understood and could perform the task before starting the experiment.

**Figure 3.**
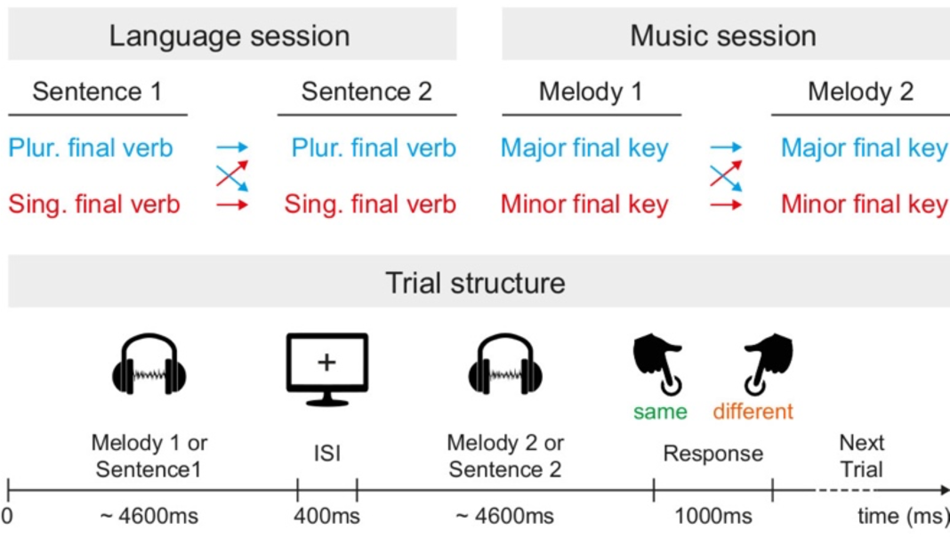
Paradigm of Experiment 1. Separately in language or music sessions, participants listened to sentence or melody pairs and had to indicate by button press whether the syntactic structure in each pair was the same or different. Sentences differed (or not) in the final verb conjugation while melodies differed (or not) in resolution of the final key (see Figure 2). Note that the trial structure is portrayed with the average stimuli duration. Plur. = plural, sing. = singular, ISI = inter-stimulus interval.

Sentences and melodies were presented with PRESENTATION (Version 15.1, www.neurobs.com) at a comfortable volume via Sony MDR-XD100 stereo headphones (Sony Corporation, Tokyo, Japan) in a quiet room. Each session was split into three blocks with 80 trials. Participants could take a one-minute break between blocks. Duration of one session was approximately 45 minutes.

### Data analysis

Data from seven participants who did not follow task instructions or performed below 50% were discarded. In the remaining 23 participants, one-sample *t*-tests verified that accuracy was significantly above chance-level (50%) in all four conditions. Reaction times (RTs) relative to the onset of the stimulus’ first critical note or word (beat/p-center 20) were extracted for correct trials only. RTs outside of two standard deviations from the mean were discarded for each participant and each condition. Analysis of variance (ANOVAs) was performed separately on accuracy (%correct) and RTs, with within-subjects factors Meter (regular, irregular) and Domain (music, language). Interactions were resolved with further *t*-tests.

## Results

Figure 4 shows accuracy and RTs in regular and irregular language and music conditions. In all conditions, accuracy was significantly above chance-level (language-regular: *t*(1,22) = 26.4, *p* < .001, *d* = 5.510; language-irregular *t*(1,22) = 26.5, *p* < .001, *d* = 5.524; music-regular: *t*(1,22) = 9.1, *p* < .001, *d* = 1.902; music-irregular *t*(1,22) = 6.9, *p* < .001, *d* = 1.442), indicating that the participants were successfully able to perform the task.

**Figure 4.**
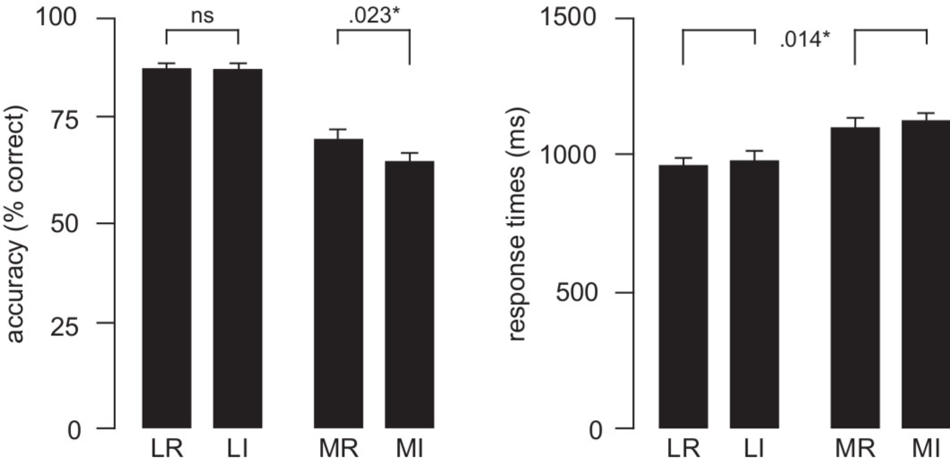
Results of Experiment 1. Syntax discrimination accuracy (left panel) and reaction times relative to the first syntactic cue (right panel) in metrically regular and irregular melodies and sentences. Error bars indicate ±1 *SEM*. L = language, M = music, R = regular, I = irregular.

Discrimination performance was significantly better in regular than irregular music. Results are summarized in Figure 4. Accuracy data showed a Meter × Domain interaction (*F*(1,22) = 4.33, *p* = .047, partial eta^2^ = .17) indicating better discrimination in regular than irregular music (paired-samples *t*-test, *t*(22) = 2.44, *p* = .023, *d* = 0.505), while no such difference was found in language (*t*(22) = 0.14, *p* = .890, *d* = 0.026). The main effect of Meter (*F*(1,22) = 4.43, *p* = .047, partial eta^2^ = .168) was likely driven by the interaction. RT data yielded a main effect of Meter (*F*(1,22) = 7.07, *p* = .014, partial eta^2^ = .243) but no Meter × Domain interaction (*F*(1,22) = 0.02, *p* = .880 partial eta^2^ = .001), indicating faster RTs in regular than irregular meter in both domains.

Additionally, responses were globally more accurate and faster in language compared to music (main effect of Domain in accuracy: *F*(1,22) = 91.12, *p* < .001, partial eta^2^ = .806, RTs: *F*(1,22) = 23.44, *p* < .001, partial eta^2^ = .516).

## Discussion

Based on previous findings (Schmidt-Kassow & Kotz, 2009; Schmuckler & Boltz, 1994), we hypothesized that syntactic resources would be comparably modulated by meter in music and language, i.e., that performance in a syntax same-different discrimination task would be better in metrically regular than irregular stimuli. This hypothesis was largely confirmed.

In language, participants discriminated syntax faster in regular compared to irregular stimuli, though accuracy did not differ across meter conditions. The RT results are in line with previous studies informing the hypothesis, supporting the notion that regular meter indeed facilitated syntax perception (e.g., Schmidt-Kassow & Kotz, 2009; Schmuckler & Boltz, 1994). Accuracy may not have aligned with RT because the task was relatively easy with performance near-ceiling.

In music, participants were both faster and more accurate in regular compared to irregular stimuli, supporting the hypothesis also in the music domain. The RT latencies of > 1100 ms suggest that participants relied on the second syntactic cue during the music task. The second cue occurred ∼400 ms after the point to which RTs were time-locked, that is, well before participants pressed the button. Moreover, from a music-theoretic perspective, it is likely that participants chose to rely on information conveyed at the strong (beat 22) rather than the weak beat (beat 20), given that syntactic information at strong beats is typically more diagnostic for musical key assignment (see *Experiment 1 introduction*).

In sum, these behavioral results support the hypothesis that syntactic resources across domains respond similarly to differences in meter. Next in Experiment 2, we investigated how meter influenced the event-related potentials (ERPs) evoked by syntactic cues in each domain.

### Experiment 2: EEG

Native listeners typically have implicit preferences for syntactic well-formedness in both language (Chomsky, 1957; Harding, Sammler, & Kotz, 2019) and music (Lerdahl & Jackendoff, 1983; Rohrmeier & Rebuschat, 2012). For example, in language, a relative clause (in italics, ‘The boss of the bus drivers *who went on strike* crossed the street’) can ambiguously modify two different nouns (boss or bus drivers) with both options being grammatical. Yet many people have an implicit preference for attaching the clause to one or the other noun (Augurzky, 2006). These preferences differ between people (Harding, Sammler, & Kotz, 2019). In music, well-formedness preference often is to resolve tension at the end of a phrase, creating closure by reiterating the tonal center of an established passage (Lerdahl & Jackendoff, 1983; Piston, 1987; Rohrmeier, 2011) as opposed to modulating to a new tonal center^4^. Among ‘native Western’ musicians, this occurs even when the new tonal center is not necessarily harmonically distant or a ‘wrong’ note, but rather closely and plausibly related (James et al., 2008).

Both single- and cross-domain studies have observed early and late negative as well as late positive ERPs in response to syntactic violations in language (Bornkessel et al., 2004; Neville et al., 1991; Osterhout & Holcomb, 1993) and music (Besson & Faïta, 1995; Featherstone et al., 2013; Koelsch et al., 2002). In a milestone cross-domain study that was influential in forming the SSIRH, Patel and colleagues (1998) showed that falsely attached relative clauses in sentences and harmonically distant chords in melodies both elicited a late positive ERP potential, a P600. The size of the P600 effect moreover increased as the falsely attached relative clauses increased processing cost, comparably in both domains. In language, the syntactic P600 effect was later shown to also be modulated by metrical grid structure in sentences (Schmidt-Kassow & Kotz, 2009), and to be significantly reduced when the meter in sentences was regular compared to irregular (Roncaglia-Denissen et al., 2013). To our knowledge, no ERP investigation has explored whether regular meter facilitates syntactic integration in music, though this intuitively should be the case, especially considering that metrical structure is part of musical syntactic well-formedness (Lerdahl & Jackendoff, 1983). Moreover, while previous studies often used syntactic violations, we explicitly use syntactic preferences in non-violated structures, to avoid confounding syntactic processing by error detection (see Slevc & Okada, 2015).

Specifically, we hypothesized that regular meter would facilitate syntactic integration in both music and language, reflected by a reduced P600 effect (e.g., Roncaglia-Denissen et al., 2013) – in other words, the regular meter should facilitate parsing, which, in turn, should make the non-preferred structure easier to integrate, thus the non-preferred ERP response should approach the preferred ERP response leading to a smaller P600 effect in regular meter.

## Materials and Methods

### Participants

Twenty-eight participants (14 female; mean age = 24.8 ± 2.5 SD) from the area around Leipzig, Germany, participated in a language and a music EEG experiment on two separate days. They were all right-handed native German speakers according to the Edinburgh handedness inventory (Oldfield, 1971) with self-reported normal hearing and no neurological history. All participants had a range of years of formal musical training as defined in Experiment 1 with a minimum of 3 and a maximum of 18 years (mean = 10.7 ± 4.3 SD). None of the participants was a pianist to avoid motor-related responses to recorded piano melodies. Participants provided written informed consent and were paid seven Euros per hour for their participation.

### Stimuli

Stimuli were the same as those described in Experiment 1.

### Tasks and procedure

A language and a music experiment took place on separate days in counterbalanced order across participants with at least two days between sessions. On both days, items were presented one by one via Sony MDR-XD100 stereo headphones (Sony Corporation, Tokyo, Japan) at a comfortable volume while participants fixated on a white cross on a black background. In the language session, the question to be answered after each item was: “Does the second phrase contain the same grammatical subject as the first phrase?” In the music session, the question was “Does the second phrase consist of the same tonal key as the first phrase?” In each session, participants first completed a computerized training of eight trials with feedback. Participants had to successfully complete the training before continuing the experiment, correctly answering the question after all condition types for stimuli and fillers.

The experiment consisted of eight six-minute blocks (60 items per block, 50% fillers) which were separated by self-paced breaks. Blocks contained items that were all regular meter or all irregular meter. The order was counterbalanced across participants. Items were pseudo-randomized such that no more than four stimuli or fillers appeared successively nor did any syntactic condition appear more than four times successively. Participants were instructed to blink between trials or during short self-paced breaks between the blocks. To avoid brain response to motor preparation and execution (e.g., from a button press), instructions during the EEG experiment were to *mentally* answer the question after each sentence or melody, and to answer with a button press only when the question was presented visually on screen after 10% of the trials. Items were separated by a 400-ms inter-stimulus interval (ISI). Preparation and testing took approximately 90 minutes per session.

### EEG recording and preprocessing

EEG was recorded with 64 Ag/AgCl electrodes placed in an elastic cap (Electro Cap Inc., Eaton, OH, USA) according to the extended 10-20 system (Sharbrough et al., 1991) using a 24 bit Brainvision QuickAmp 72 amplifier (Brain Products GmbH, Gilching, Germany). Sampling rate was 500 Hz. Impedances were kept below 5 kΩ throughout the experiment. Eye movements were monitored by bipolar horizontal and vertical electrooculograms (EOG) recorded from electrodes placed beneath the canthus of both eyes as well as above and below the right eye. Additionally, electrodes were placed on the left and right mastoid bones (M1 and M2) and a ground was placed on the sternum. The reference during recording was M1.

Offline, EEG data were re-referenced to linked mastoids and high-pass filtered at 0.4 Hz to remove excessive drifts (−3dB cutoff, fir, 3333 points, Blackman window) with EEProbe (ANT neuro, Enschede, NL). The high-pass filter also served as a baseline filter. Each trial was determined by the length of the corresponding stimulus and subsequent ISI (approximately 5.2 seconds), and all further preprocessing steps were applied on these epoched data. Epochs were automatically removed when they contained large artifacts, i.e., whenever the standard deviation of the signal in a 200 ms sliding window exceeded 20µV. With FieldTrip (Oostenveld et al., 2010), ICA was then used to correct blinks and eye-movements. Five frontal electrodes (FPz, FP1, FP2, AF3, AF7) that were a source of noise in several participants were removed (in the language data only). In one participant, one electrode within the regions of interest (described below) was removed and subsequently interpolated. Finally, cleaned data were visually inspected and trials with remaining artifacts were discarded. Participants with more than 20 rejected trials per condition (i.e., less than 40 trials) were excluded from further analysis. One participant was excluded for not correctly completing the task during recording. The final group size was 23 participants.

### ERP analysis

Further ERP analyses were conducted in smaller epochs from −600 ms to 1000ms, time-locked to the onset of the critical final morpheme of the verb *besuch**te/n*** in sentences or the onset of the final note in melodies. The epochs included the end of the respective stimulus and the following ISI. In both language and music sessions, trials were averaged for preferred and nonpreferred syntactic constructions, separately for regular and irregular meter. Syntactic preference was determined on a per-participant basis in a post-hoc questionnaire. That is, at the end of the language session, participants were asked whether they preferred to attach the pronoun to the subject (high attachment) or the possessor (low attachment). This coding method was previously validated in auditory German relative clause attachment (Harding, Sammler, & Kotz, 2019). In the music session, participants indicated whether they preferred the original as opposed to the modulated key. The condition that participants listed as “preferred” was then coded as such in the ERP analysis. In music, all participants preferred the original major endings, consistent with previously reported Western tonal preferences (Nieminen et al., 2012; Parncutt, 2012). In language, 13 participants preferred high attachment, 5 preferred low attachment, and 5 had no preference. Their coding was assigned the default of low-attachment preference, based on previous work with auditory German sentences with this particular type of Genitive possessor relative clause attachment (Augurzky, 2006). Low attachment was also the majority attachment preference in Harding, Sammler and Kotz (2019).

To identify relevant time windows of the P600, preferred- and non-preferred syntax conditions (collapsed across meter type) were first compared using cluster-based permutations using the Monte Carlo method with 1000 iterations and 0.05 alpha threshold (Oostenveld et al., 2010). In language, a positive cluster with broad scalp distribution was found between 738ms and 834ms, time-locked to the syntactically critical verb morpheme “-ten/-te”. This positivity was visually inspected and taken as a P600 effect, despite a slightly later latency than 600ms, consistent with previous auditory domain studies (Canette et al., 2020; Patel et al., 1998). In music, a significant positive cluster was found between 656ms and 888ms, time-locked to the final note, that after visual inspection was labeled a P600 (Besson & Faïta, 1995). The identified time-windows for the hypothesized-for positive clusters were used to investigate the effects of metrical regularity on syntactic processing in midline and lateral regions of interest (ROI): Midline ROIs were anterior (AFz, Fz, FCz, Cz) and posterior (CPz, Pz, POz, Oz). The lateral ROIs were sectioned as follows: left anterior (F7, F5, F3, FT7, FC5, FC3), left middle (T7, C5, C3, TP7, CP5, CP3), left posterior (P7, P5, P3, PO7, PO3, O1), right anterior (F4, F6, F8, FC4, FC6, FT8), right middle (C4, C6, T8, CP4, CP6, TP8), right posterior (P4, P6, P8, PO4, PO8, O2). Averaged-over-ROI, averaged-over-time-windows amplitude values were used to conduct 1) a midline ANOVA with repeated-measures factors Syntax (preferred, non-preferred), Meter (regular, irregular), and AntPost (anterior midline, posterior midline) and 2) a lateral ANOVA with repeated-measures factors Syntax (preferred, non-preferred), Meter (regular, irregular), Hemisphere (left, right), and AntPost (anterior, middle, posterior) in each domain.

## Results

### Language

Both the midline and the lateral ANOVAs showed significant main effects of Syntax, but no interactions with topographical factors Hemisphere or AntPost, indicating that the P600 effect had a broad scalp distribution (see top panels in Figure 5 and Table 1 for statistical values). This P600 effect was significantly larger in the regular than the irregular meter condition as indicated by a significant Syntax × Meter interaction in the midline ANOVA. The interaction was driven by differences between regular and irregular meter occurring in the preferred syntax condition only (see Figure 6; regular/preferred vs. irregular/preferred: *F*(1,22) = 10.925, *p* = .003, partial eta^2^ = .332; regular/non-preferred vs. irregular/non-preferred: *F*(1,22) = 0.033 *p* = .857, partial eta^2^ = .002). Similarly, a marginal Syntax × Meter × Antpost interaction was found in the lateral ANOVA. When resolved by step-down ANOVAs with factors Syntax, Meter, and Hemisphere in anterior, middle and posterior ROIs, it revealed a near-significant Syntax × Meter interaction in the middle ROI (*F*(1,22) = 4.153, *p* = .054, partial eta^2^ = .159), but not the anterior or posterior ROIs (*F*s < 1.812, *p*s > .192, partial eta^2^ < .076). There were no other significant main effects or interactions (see Table 1 for all statistical values).

**Figure 5.**
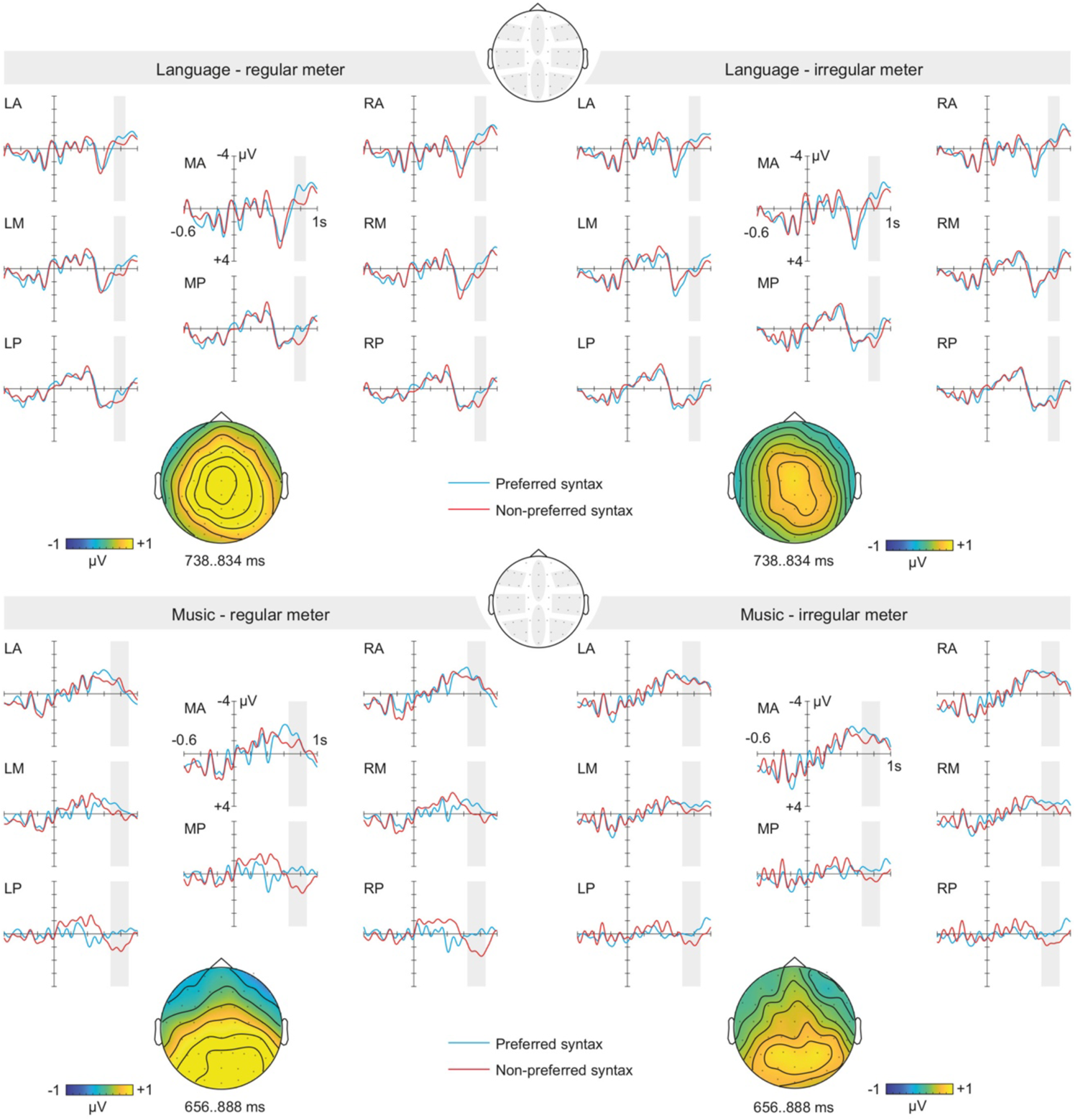
*Language, upper panel.* A broadly centrally-distributed P600 effect time-locked to the final verb conjugation was reduced in the irregular meter condition. *Music, lower panel.* A posteriorly distributed P600 effect time-locked to the final note showed a trend to be reduced in the irregular meter condition.

**Figure 6.**
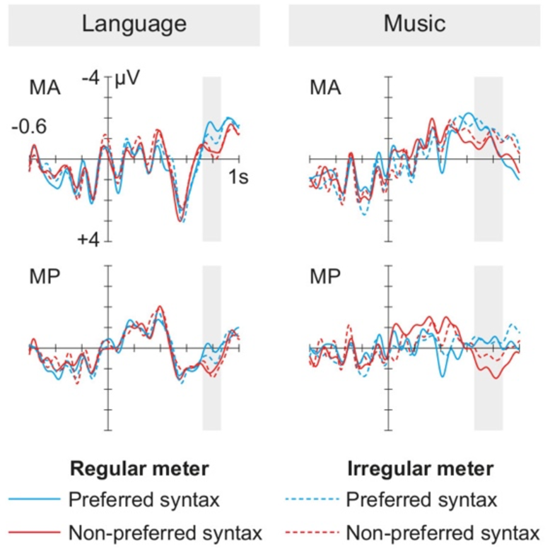
Illustration of the Syntax × Meter interaction in language and music. In language, the reduced P600 effect in irregular (dashed lines) compared to regular meter (solid lines) was explained by a modulation of the preferred condition (blue lines). In music, the reduced P600 effect in posterior areas was explained by a negative shift of the non-preferred condition (red dashed line). Thus, while across domains the overall impact of irregular meter was a reduced P600 effect, the meter interaction with syntactic-integration-related cognitive processes seems to be qualitatively different.

**Table 1.** Results of the repeated-measures ANOVAs conducted on the ERPs.

|  | Language P600 |  |  | Music P600 |  |  |
| --- | --- | --- | --- | --- | --- | --- |
| | *F*(1,22) | *p*-value | $\eta_p^2$ | *F*(1,22) | *p*-value | $\eta_p^2$ |
| *Midline ANOVA* |  |  |  |  |  |  |
| S | **22.827** | **.001** | **.509** | **14.955** | **.001** | **.405** |
| M | **3.805** | **.064** | **.147** | **6.829** | **.016** | **.237** |
| S × M | **4.430** | **.047** | **.168** | 1.088 | .308 | .047 |
| S × AP | 0.347 | .562 | .016 | **19.863** | **< .001** | **.474** |
| M × AP | 0.191 | .667 | .009 | 0.371 | .548 | .017 |
| S × M × AP | 0.370 | .549 | .017 | **3.807** | **.064** | **.148** |
| *Lateral ANOVA* |  |  |  |  |  |  |
| S | **16.558** | **.001** | **.429** | **13.389** | **.001** | **.378** |
| M | 2.918 | .102 | .117 | **7.308** | **.013** | **.249** |
| S × M | 2.885 | .104 | .116 | 0.880 | .358 | .038 |
| S × AP | 2.083 | .150 | .166 | **14.799** | **<.001** | **.585** |
| M × AP | 0.129 | .879 | .012 | 0.254 | .778 | .024 |
| S × M × AP | **2.095** | **.073** | **.220** | **2.614** | **.097** | **.199** |
| S × H | 1.243 | .277 | .053 | 1.485 | .236 | .063 |
| M × H | 0.864 | .363 | .038 | **4.158** | **.054** | **.159** |
| S × M × H | 1.507 | .233 | .064 | 1.001 | .328 | .044 |
| S × AP × H | 0.434 | .654 | .040 | 2.045 | .154 | .163 |
| M × AP × H | 0.654 | .530 | .059 | 1.777 | .194 | .145 |
| S × M × AP × H | 0.574 | .572 | .052 | 0.549 | .586 | .050 |

Significant effects in bold, marginal effects in italicized bold. S: Syntax; M: Meter; AP: Anterior-Posterior; H: Hemisphere.

### Music

In the P600 time-window, both midline and lateral ANOVAs showed significant Syntax main effects and Syntax × AntPost interactions, indicating that the P600 had a central-posterior distribution (see lower panels in Figure 5 and Table 1). The Syntax × Meter interaction was not significant, however, considering the strong hypothesis that meter modulates syntactic integration, we followed up a marginal Syntax × Meter × AntPost interaction in midline and lateral ROIs. Results were consistent with a Syntax × Meter interaction in posterior regions where the P600 was strongest. In these regions, the P600 effect was larger in the regular than the irregular meter condition (see Figure 5, bottom panel), that is, the interaction approached significance both in the posterior lateral ANOVA (*F*(1,22) = 4.12, *p* = .055, partial eta^2^ = .158) and the posterior midline ANOVA (*F*(1,22) = 3.50, *p* = .076, partial eta^2^ = .138; all other ROIs *F*s < 0.90, *p*s > .3). Notably, these interactions were driven by differences between regular and irregular meter occurring in the *non-*preferred syntax condition only (see Figure 6; regular/preferred vs. irregular/preferred in the posterior midline ROI: *t*(22) = 0.141, *p* = .889, Cohen’s *d* = .029; regular/non-preferred vs. irregular/non-preferred: *t*(22) = 2.308, *p* = .031, Cohen’s *d* = .481; regular/preferred vs. irregular/preferred in the posterior lateral ROIs: F(1,22) = 0.41, p = .841, partial eta2 = .002; regular/non-preferred vs. irregular/non-preferred: F(1,22) = 4.135, p = .054, partial eta2 = .158).

Moreover, the ANOVA showed a significant main effect of Meter indicating that amplitudes were generally larger in the regular meter as opposed to irregular meter condition. A near significant Meter × Hemisphere interaction in the lateral ANOVA indicated that this difference was slightly stronger in the right hemisphere (left: *F*(1,22) = 4.136, *p* = .054, partial eta^2^ = .158; right: *F*(1,22) = 9.42, *p* = .006, partial eta^2^ = .300). There were no other significant effects for the P600 (see Table 1 for detailed statistical values).

## Discussion

Supporting our hypothesis and consistent with the literature, Experiment 2 revealed a P600 elicited by non-preferred syntactic structures in both language (Bornkessel et al., 2004; Neville et al., 1991; Osterhout & Holcomb, 1993) and music (Besson & Faïta, 1995; Featherstone et al., 2013; Koelsch et al., 2002). However, while a regular meter seemed to facilitate the behavioral discrimination of syntactic constructions in Experiment 1, the role of the meter was more nuanced in Experiment 2. We had hypothesized that, in line with Experiment 1 and previous results from the language domain (Roncaglia-Denissen et al., 2013), regular meter should facilitate the integration of non-preferred syntactic constructions, and that this would be reflected in ERP waveforms from the non-preferred condition approaching the preferred amplitude in regular sentences, leading to a reduction of the P600 effect. However, the ERPs rather reflected that meter reduced the P600 effect in *ir*regular conditions across domains, and further, that the interaction with syntactic integration was qualitatively different across domains.

In language, the P600 effect was smaller in metrically irregular compared to regular sentences, because preferred-syntax potentials in the regular meter condition modulated towards resembling non-preferred potentials in the irregular meter condition (dashed blue line in Figure 6). Moreover, metrical regularity seems not to have affected the integration of non-preferred syntactic structure, as these non-preferred potentials remained largely unchanged across meter conditions (red lines in Figure 6). This finding is contrary to our hypothesis that regular meter should facilitate the integration of non-preferred syntactic structures. This departure from previous results in non-musicians (Roncaglia-Denissen et al., 2013) may be due to the different testing population. Indeed musicians are more sensitive to speech meter than non-musicians (Magne et al., 2007; Marie et al., 2011). Thus, musicians—the target group of the present study necessary to compare the processing of syntactic complexity in music—may have quickly adapted to and utilized the regular meter as a template for syntactic parsing and were thus affected by its disruption in the irregular meter condition.

In the music domain as well, the P600 effect was reduced in metrically *ir*regular compared to regular melodies, contrary to our hypothesis that a regular meter would facilitate syntactic integration by reducing the effort necessary to integrate the non-preferred syntax. Interestingly, the reduction of the P600 effect was qualitatively different in music compared to language: in music, the potentials for *non*-preferred (rather than preferred) syntax structure were reduced in the irregular meter context (red dashed line in Figure 6), suggesting that there was little cost to integrate the non-preferred structure. Considering that irregular metrical structures are less common in Western classical music, it is not likely that the irregular meter *facilitated* syntactic integration. Rather, a reasonable explanation is that a regular meter is a prerequisite to syntactic well-formedness in classical music, and that if the metrical context is unusual or unexpected, this may suspend the expectations for Western-tonal well-formedness. For example, in 20^th^ century musical traditions, irregular rhythms and atonality often occur in tandem. Thus, perhaps the musician participants did not register the modulation to a relative minor key as less well-formed when the metrical context was irregular. Considering that meter has a defining role in the syntactic structure of music (Lerdahl & Jackendoff, 1983), perhaps what we see is that syntactic integration for typically Western classical tonality is framed by metrical well-formedness and breaks down when meter is irregular.

Considering that the perception of both syntax and meter is subject to individual differences in working memory (Bornkessel et al., 2004), temporal perception (Skoe & Kraus, 2010), and musical ability (Geiser et al., 2010; Harding, Sammler, Henry, et al., 2019; Marie et al., 2011), looking at these aspects of cognition may explain some of the experimental findings (Experiment 1 & 2). To what extent these individual differences can account for the present behavioral and electrophysiological results was further addressed in Experiment 3.

### Experiment 3: Individual differences

The final leg of the current investigation developed continuous individual difference metrics that were then used to explain variance in the influence of regular meter on syntax processing. We included measures of temporal perception, musical ability, and working memory because each has documented links to syntax processing across domains.

In language, temporal perception has been linked to the fine-grained perception of syllable features such as phonemes (Goswami, 2011; Tierney & Kraus, 2014), which are important prerequisites to discriminating syntactic inflections (such as ‘-ten’/’-te’) that can determine syntactic structure. Increased musical ability is also linked to improved speech encoding and categorical perception of speech (Mankel et al., 2020; Mankel & Bidelman, 2018), both steps necessary for syntactic discrimination. Working memory has been associated with the ability to build and maintain complex syntactic structures (e.g., Nieuwland & Van Berkum, 2006), including the ability to better inhibit information that competes with preferred meanings of ambiguous lexical items or syntactic structures (Bornkessel et al., 2004; Gunter et al., 2003; Meyer et al., 2013).

In music, discriminating rhythms with temporal differences has been correlated with discriminating melodies differing in syntactic construction (Wallentin et al., 2010), and temporal change detection mechanisms are suggested to interact with processing harmonic tonality of musical syntax (Lebrun-Guillaud & Tillmann, 2007). Musical ability is linked to superior perception of tones (Wayman et al., 1992) and syntactic-harmonic structure (e.g., Besson & Faïta, 1995). The role of working memory in music was previously demonstrated by poorer task performance when musical syntactic errors are introduced during syntax structure-building in visual language (Fedorenko et al., 2009; Fiveash & Pammer, 2014). Considering this evidence, we tested these auditory perceptual abilities with the aim to see to what extent they may play a role in how meter interacts with syntactic-discrimination and integration across language and music domains.

In the literature, musical expertise (i.e., being a musician) often overlaps with individual differences in temporal perception, musical ability, and working memory. Musicians have been reported to have, as a group, higher verbal working memory capacity than non-musicians (Chan et al., 1998; Ho et al., 2003), and higher auditory working memory is also linked to better temporal perception (Tzounopoulos & Kraus, 2009). While musical ability has clear links to degrees of musical expertise (Wallentin et al., 2010), sophisticated musical ability is also found in non-musician populations (so-called “musical sleepers”, Mankel & Bidelman, 2018). Thus, with a secondary aim to explore the advantage of thorough diagnostic testing compared with traditional musical expertise categorization, we assessed individual difference metrics according to musician and non-musician grouping, age of musical training onset and hours of weekly practice.

We administered seven auditory diagnostic tests to 88 participants with varied musical training and used principal component analysis (PCA) to identify latent variables underlying their performance. The diagnostic tests were the forward digit span and backward digit span (Tewes, 1994), non-word repetition (Mottier, 1951), anisochrony detection (Dalla Bella et al., 2017), and rhythm and melody subtests of the Musical Ear test (Wallentin et al., 2010). The PCA resulted in two streamlined components reflecting Verbal Working Memory (VWM) abilities and acuity in Time and Pitch Discrimination (TPD). To assess the extent to which these components play a role in cross-domain syntax discrimination and syntactic integration, these individual difference metrics were entered into correlations with the syntax discrimination measures from Experiment 1 and ERP P600 effects from Experiment 2, respectively. Additionally, the metrics were assessed according to musical expertise.

## Materials and methods

### Participants

Eighty-eight native German speakers (50 female, mean age ± *SD*: 25 ± 7 years) were recruited from the participant database of the Max Planck Institute for Human Cognitive and Brain Sciences, Leipzig, Germany. Thirty of them were self-reported non-musicians and 58 were self-reported musicians with a broad range of musical training. The average age of training onset among musicians was 11.1 years (*SD* = 4.8) and average number of practice hours per week was 5.38 (*SD* = 4.25) at the time of testing. All participants from Experiment 1 & 2 were included in this sample. Participants gave written informed consent prior to the experiment and received 7 Euros per hour for their participation. The study was approved by the ethics committee of the University of Leipzig and carried out following the guidelines of the declaration of Helsinki.

### Diagnostic tests

#### Musical Ear Test (MET)

The Musical Ear Test (MET; Wallentin et al., 2010) tests melody and rhythm discrimination in two separate subtests, each containing 52 trials. In each trial, participants judge whether two short musical phrases are the same or different. The melody subtest presents melodies of 3-8 tones at 100 bpm that differ in one tone (or not). The rhythm subtest presents beat sequences with 4-11 wood-block sounds that contain one rhythmic change (or not). Items vary greatly in difficulty avoiding floor and ceiling effects in participants with no or high musical training. We registered the total correct answers per subtest per participant (score range: 0-52).

#### Forward and backward digit span (FDS & BDS)

Forward and backward auditory digit span (FDS & BDS) tests were used from the Wechsler Intelligence Scale (Tewes, 1994) as widely accepted measures of working memory capacity. In these tasks, participants are asked to repeat a spoken series of digits, either forwards (FDS) or backwards (BDS). The number of digits incrementally increases across the test, from two (BDS) or three (FDS) to nine digits per series. We registered the length of the longest correctly remembered series for FDS and BDS (score range FDS: 3-9; BDS: 2-9).

#### Non-word repetition

The nonword repetition test (Mottier, 1951) measures phonological working memory capacity and the ability to discriminate phonemes. In this test, participants are asked to repeat spoken pseudo-words with incrementally increasing number of syllables, beginning with two syllables (e.g., ‘re-la’) up to eleven syllables (e.g., ‘su-wa-mu-fo-bo-ga-ku-po-mi-so-ti’). We registered the highest correctly repeated syllable count (score range: 2-11).

#### Modified listening span

The modified listening span (Daneman & Carpenter, 1980) was designed to index the trade-off between storage and manipulation in working memory, specifically as it pertains to language comprehension. Participants are presented with spoken sentences (e.g., ‘All coats are brown.’) in blocks of three to seven sentences. After each sentence per block, they have to first repeat the last words of all sentences heard before in that block, before answering whether the statement of the current sentence was true or false. We registered the length of the longest block correctly mastered by participants (score range: 3-7).

#### Anisochrony detection

The anisochrony detection test (adapted from Dalla Bella et al., 2017) estimates participants’ temporal discrimination threshold using a maximum likelihood procedure. The task includes five tones (frequency = 1047 Hz, duration = 150 ms) with an interval of 600 ms (onset to onset). The fourth tone is adaptively displaced up to 180 ms earlier depending on the participant’s judgment whether the series was regular or not. Randomly inserted ‘catch trials’ without displacement allow controlling participants’ attention to the task. The just noticeable displacement (in ms) was estimated three times per participant, in three blocks containing sixteen adaptive 5-tone sequences each. The three threshold values were averaged to obtain the participant’s final threshold. Blocks in which (i) more than 30% of the ‘catch trials’ were erroneously judged as irregular, or in which (ii) the threshold estimate changed by more than 15% over the last 10 non-catch trials (meaning the maximum likelihood algorithm was not converging) were discarded. An average of 2.86 blocks were entered into the analysis per participant.

#### General Procedure

Computerized versions of all diagnostic tests were administered via headphones at a comfortable volume in a quiet room. Tests that contained words used recordings spoken by professionally trained, female native German speakers. Instructions were either written or spoken aloud by the test administrator. Correct-hand response was counterbalanced across participants but kept consistent for tests within participants, and the order of diagnostic test presentation was pseudo-randomized. The total amount of time needed to complete all tests was approximately 1.5 hours.

### Data analyses

#### Principal Component Analysis (PCA)

The seven diagnostic scores (MET-melody, MET-rhythm, FDS, BDS, non-word repetition, modified listening span, anisochrony threshold) were entered into a principal component analysis (PCA) to identify and compute composite scores of underlying latent variables. The analysis was performed with SPSS Version 20 (IBM, Armonk, NY). Data from two participants were excluded because their anisochrony thresholds were identified as extreme outliers, that is, they were more than 3 times the interquartile range lower (higher) than the first (third) quartile of the group. The final sample size of 86 and the seven diagnostic test scores fulfilled the assumptions for PCA (Field, 2005): First, six of the seven test scores correlated at least *r* = .300 with at least one other item, suggesting reasonable propensity to cluster into components. Second, the Kaiser-Meyer-Olkin measure of sampling adequacy was .67, above the commonly recommended value of .60, and Bartlett’s test of sphericity was significant (*F*(2,21) = 84.14, *p* < .001). The diagonals of the anti-image correlation matrix were also all over .50, and the determinant was greater than 0.00001 (.358), indicating no problems with multicollinearity. An oblimin rotation provided the best-defined factor structure. The pattern matrix was consulted to interpret the unique contribution of each test score to its principal component; all test scores had primary loadings over .5, and cross-loadings were all below .2. Finally, the communalities were all above .30, further confirming that each item shared some common variance with other items. After computation of the PCA, consistency of the test scores per principal component was estimated using Cronbach’s alpha. Stability of principal component loadings was tested using a jackknife procedure, whereby each participant was removed and the principal component loadings re-estimated with the remaining 85 participants.

#### Musical expertise

Although we focussed on musical ability and not on musical expertise per se (see Mankel & Bidelman, 2018), we aimed to see whether the principal component scores (PC-scores) were impacted by musical expertise comparably to previous studies. PC-scores were first compared between musicians and non-musicians (e.g., Ho et al., 2003). Two self-reported non-musicians were excluded from this analysis because they had received extracurricular musical training during childhood. We also excluded four self-reported musicians who no longer were actively playing their instrument, resulting in a final sample of 26 non-musicians (mean age = 26.3 years, *SD* = 3.1, 11 female) and 54 musicians (mean age = 23.4 years, *SD* = 2.5, 31 female). Two-sample *t*-tests were performed with the hypothesis that musicians would have higher scores than non-musicians (Rammsayer & Altenmüller, 2006; Tzounopoulos & Kraus, 2009). Within the musician group we calculated Spearman correlations between the PC-scores, training onset (e.g., Wong et al., 2007), and practice hours (e.g., Wallentin et al., 2010).

#### Correlations with behavioral and ERP findings

Finally, Spearman correlations were conducted to see whether the behavioral results from the syntax discrimination task (Experiment 1) or the ERPs in the syntactic integration task (Experiment 2) were statistically related to participants’ PC-scores in VWM and TPD. All analyses were Bonferroni-corrected for multiple comparisons.

## Results

### Principal component analysis

Initial eigenvalues indicated that the first two principal components explained 33% and 19% of the variance, respectively. The component labels were assigned according to the common cognitive functions responsible for their respective tasks: Verbal Working Memory (VWM) and Time and Pitch Discrimination (TPD). The pattern matrix is presented in Table 2. The third through seventh components had eigenvalues below one and were therefore not further considered. The two-component solution, which explained 52% of the variance, was accepted because of theoretical support for clustering of verbal working memory and temporal and pitch discrimination on separate components, as well as the insufficient number of primary loadings and difficulty of interpreting subsequent components. Although Cronbach’s alpha indicated moderate consistency for principal component 1 (.66 for four items), and poor consistency for component 2 (.55 for three items), a jackknife procedure showed that the two components remained constant and retained their individual diagnostic test contributions across 86 reiterations of the PCA. Therefore, and because all other assumptions of the PCA were met, the internal consistency of the two-principal component solution was deemed sufficient.

**Table 2.** Principal component loadings and communalities based on the PCA with oblimin rotation for the seven diagnostic test scores.

|  | VWM | TPD | Communality |
| --- | --- | --- | --- |
| Backward digit span | .80 |  | .62 |
| Forward digit span | .79 |  | .60 |
| Modified listening span | .59 |  | .41 |
| Non-word repetition | .58 |  | .42 |
| MET melody subtest |  | .74 | .59 |
| Anisochrony detection |  | .70 | .55 |
| MET rhythm subtest |  | .68 | .47 |

VWM = Verbal Working Memory (component 1), TPD = Time and Pitch Discrimination (component 2). Component loadings < .20 are suppressed.

For further analyses, one standardized composite score was extracted per participant per principal component (VWM or TPD), which combined all tests that loaded onto that principal component. Higher scores indicate greater aptitude in the cognitive functions underlying VWM and TPD.

### Musical expertise

The left panel in Figure 7 shows that musicians scored higher in TPD than non-musicians (mean ± *SD* in musicians: 0.38 ± 0.85; non-musicians: −0.70 ± 0.83; *t*(78) = 5.39, *p* < .001, *d* = 1.30), while the two groups did not differ in VWM (musicians: 0.02 ± 1.02; non-musicians: −0.11 ± 0.99; *t*(78)= 0.56, *p* = .58, *d* = 0.13). Within the musician group, practice hours in the year preceding the experiment correlated with TPD (Spearman’s *rho* = .32, *p* = .018), but not VWM (*rho* = .02, *p* = .892). No significant correlations were found between age at training onset and TPD (*rho* = .01, *p* = .964) or VWM (*rho* = −.04, *p* = .766).

**Figure 7.**
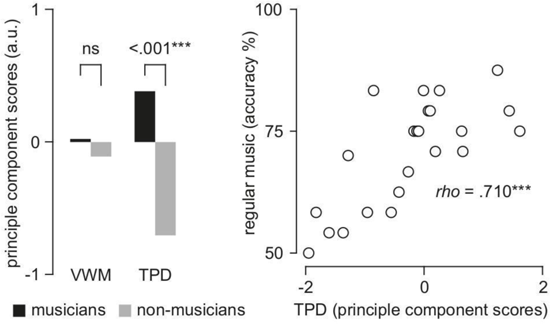
Relationship between musical expertise, working memory, temporal perception and syntax processing. *Left panel:* Principal component scores assessed according to musical expertise. Time and Pitch Discrimination (TPD) was significantly higher in musicians than non-musicians, while no expertise effects were found for Verbal Working Memory (VWM). *Right panel:* TPD scores correlated with participants’ accuracies while discriminating syntax of metrically regular music in Experiment 1.

### Correlations with behavioral and ERP findings

#### Behavior (Experiment 1)

Accuracies and mean RTs of participants in the syntax discrimination task (Experiment 1) were entered into correlations to test whether there was a statistical relationship with participants’ scores in VWM and in TPD (accuracy and RT in regular-music, irregular-music, regular-language, and irregular-language conditions; alpha was Bonferroni-corrected to a significance threshold of *p* < .006). Participants’ syntax discrimination accuracy in the regular-music condition significantly correlated with TPD, such that the greater the TPD score, the greater the accuracy discriminating major from minor syntax in regular-meter melodies. Results are displayed in Table 3. No other significant links were found between participants’ syntax discrimination performance and cognitive factors.

**Table 3.** Correlations between PC-scores and syntax discrimination performance

|  | VWM |  | TPD |  |
| --- | --- | --- | --- | --- |
|  | *rho* | *p*-value | *rho* | *p*-value |
| *Accuracy* |  |  |  |  |
| LR | .240 | .269 | .159 | .469 |
| LI | .065 | .770 | -.032 | .886 |
| **MR** | .340 | .112 | **.709** | **< .001** |
| MI | .451 | .031 | .360 | .092 |
| *Response times* |  |  |  |  |
| LR | -.076 | .730 | -.310 | .150 |
| LI | -.130 | .553 | -.378 | .075 |
| MR | -.112 | .612 | -.133 | .544 |
| MI | .024 | .914 | -.011 | .961 |

Significant effects in bold (Bonferroni-corrected).

#### ERPs (Experiment 2)

Correlations were used to see whether syntax ERP effects (non-preferred minus preferred) were associated with VWM and TPD for regular and irregular meter conditions in the mid-posterior ROI where the P600 effect was largest. In total there were 4 correlations for each cognitive factor (see Table 4). Alpha was Bonferroni-corrected for eight comparisons (\**p* < .006). None of the correlations reached significance.

**Table 4.** Correlations between PC-scores and amplitude of the P600 effect in the mid-posterior

|  | VWM |  | TPD |  |
| --- | --- | --- | --- | --- |
|  | *rho* | *p*-value | *rho* | *p*-value |
| *Mid-posterior ROI* |  |  |  |  |
| LR | .104 | .638 | -.010 | .964 |
| LI | .163 | .457 | .054 | .805 |
| MR | .011 | .961 | .150 | .494 |
| MI | .262 | .227 | .404 | .056 |

## Discussion

Experiment 3 successfully classified cognitive abilities shown in previous studies to be relevant for cross-domain syntax processing, gaining two scores that were then used to assess individual differences in syntax processing from Experiments 1 and 2. A principal component analysis assessed individual differences in musical ability, temporal perception, and working memory. Principle components were named Verbal Working Memory (VWM) and Time and Pitch Discrimination (TPD), appropriate to the cognitive functions associated with tasks in the respective diagnostic tests found in each principal component.

The advantage of using a continuous cognitive factor as a metric is that, on the one hand, splitting analysis by groups is statistically less appealing than using a continuous metric (MacCallum et al., 2002). On the other hand, grouping according to a background of musical training has the potential to include participants with good musical ability in non-musician groups (Mankel & Bidelman, 2018). While the cognitive factors were devised to look beyond musician-non-musician groupings, we still, however, evaluated the factors according to expertise to verify whether the factors were consistent with the musicianship literature. Indeed, musicians had a higher TPD mean score than non-musicians, which is in line with previous findings that musical expertise impacts melodic contour, rhythm- and real-time perception of temporal information (Rammsayer & Altenmüller, 2006; Wallentin et al., 2010). The correlation with weekly practice hours (but not age of training onset) suggests that perhaps these abilities might be sensitive to short-term fluctuations in training or musical exposure.

Verbal working memory did not differ between musical expertise metrics or grouping. This finding contradicts previous observations in the literature that reported higher verbal working memory in musicians compared to non-musicians (Chan et al., 1998; Franklin et al., 2008; Ho et al., 2003; Tzounopoulos & Kraus, 2009). This may be due to the current criteria for musicians including participants who started musical training until late adolescence and without regard to a history of consistent practice, compared to more stringent criteria regarding training age-of-onset and uninterrupted weekly practice in previous studies (Lotze et al., 2003; Strait et al., 2012). Thus perhaps the musicians as a group missed some critical period for plasticity (e.g., Pantev et al., 1998; Schlaug, 2001), which would have offered better functional networks associated with verbal working memory. Alternatively, grouping according to a background of musical training has the potential to capture other factors not related to musical ability such as socioeconomic status (Hoffman, 2013), personality, or an innate predisposition to take music lessons (Corrigall et al., 2013). Thus, previous findings may have reflected increased verbal working memory abilities linked to children growing up in households where parents could send their children to music lessons from an early age without interruption, or that children with high verbal working memory may generally succeed in extra-curricular activities if they e.g., can spend less time on other schoolwork. Future research is warranted to more closely investigate verbal working memory differences in musicians and non-musicians. Accuracy of syntactic discrimination in regular-meter melodies was found to correlate with TPD, such that performance improved as TPD increased. This is intuitive when considering that people who are better at perceiving “time” benefit more from regular meter in syntax perception. TPD, but not VWM, was furthermore linked to musical expertise metrics, suggesting that it may underlie musician-non-musician group differences reported in other studies. The cognitive factors did not significantly correlate with any other experimental findings.

### General Discussion

The current set of studies aimed at assessing comparability of syntactic processing in music and language while considering different metrical composition in both domains in addition to individual differences in auditory perception and cognitive abilities. In three experiments, results show that syntactic processing interacts with meter in both music and language, albeit differently, and that syntax performance in the music domain was associated with individual differences in auditory temporal perception.

Novel stimuli, comprising syntactic complexities, elicited a P600 effect in each domain. As the syntactic complexities tested preference rather than error-detection and repair, the P600 findings, thus, appear to reflect syntactic integration in both domains (Patel, 2003) rather than error-detection and repair or cognitive control (Slevc & Okada, 2015). Behavioral results confirmed a facilitation effect of regular meter on syntactic discrimination in both domains (Experiment 1), and meter also modulated the P600 effect similarly across domains (Experiment 2). At first glance this seems to reflect a similar impact of meter on syntax, however, the denoted details suggest a more nuanced picture.

We found that regular meter facilitated the behavioral syntax discrimination in both domains, in a group including both musicians and non-musicians (Experiment 1). This is in line with multiple studies suggesting that (temporally) regular meter can facilitate language and music syntax processing. For example, Schmidt-Kassow and Kotz (2008) previously showed that syntactic comprehension in language is influenced by the temporal predictability of salient lexical items, and that the metrical identity of words was a necessary perceptual landmark for syntactic reanalysis (Schmidt-Kassow & Kotz, 2009). Roncaglia-Denissen et al. (2013) further showed syntactic integration of correct, non-preferred structure to be facilitated when presented in a regular metrical context. In the music domain, the present paradigm seems to be the first to vary meter independently from the tonal components of syntax, though previously, temporal jitter in meter disrupted syntactic comprehension (Nittono et al., 2000) or modulated the judgment of harmonic appropriateness of chords (Bigand et al., 1999; Schmuckler & Boltz, 1994).

However, ERPs measured in musicians in a syntactic comprehension task revealed a larger (rather than the expected smaller) P600 effect for non-preferred syntactic structure in a regular meter context (Experiment 2). This seems to contradict the results of Experiment 1 as well as the literature, because a larger P600 effect may suggest more effortful processing as opposed to facilitated processing. In language, a possible reason for this deviation of the P600 effect from the literature is that, the previous ERP study, upon which we based the present language P600 hypothesis, tested non-musicians (Roncaglia-Denissen et al., 2013), whose results may not generalize to musicians. In music, an emerging explanation for the direction of the P600 modulation is that a regular meter is actually a prerequisite for syntactic structure building and not just a facilitating component for syntactic integration. Without clear metrical boundaries that usually assign harmonic boundaries (Bigand et al., 1999; Boltz, 1993) there may be no awareness of a harmonic complexity that would tax syntactic integration processes in the first place. These possibilities will be discussed in turn.

### Language

Prior results suggest that a regular meter facilitates syntactic integration in non-musicians, marked by a reduced P600 effect (Roncaglia-Denissen et al., 2013). While we replicated a syntax-meter interaction in the current study with musicians, we found it in the opposite direction—the P600 effect was reduced in the *ir*regular (rather than regular) meter condition. The previous design was similar to the current one in that locally ambiguous relative clauses were presented in regular and irregular meter conditions, and the meter was constructed by varying the number of syllables in content words (e.g., “Ghent” vs. “Brussels”). Thus, a likely source of the discrepancy is the testing population. Considering that musicians have been observed to be more sensitive to speech meter than non-musicians (Magne et al., 2007; Marie et al., 2011), here musician participants may have immediately recognized the regular metrical pattern and utilized it to inform syntactic parsing. This syntactic processing may have, in turn, been disrupted when encountering irregular meter. The disruption of syntactic processing for *violated* meter has precedence when looking closely at the literature. Schmidt-Kassow and Kotz (2009) investigated the interaction of meter and syntax perception in non-musicians, and found P600 effects in three conditions: violated syntax, violated meter, and a combined syntax-meter violation. Violated syntax was created by an ungrammatical verb conjugation. Violated meter was created by stressing the wrong syllable in words (e.g., “visit” pronounced as vi-’SIT instead of ‘VIS-it). Notably, the P600 effect evoked by the combined violation was under-additive, i.e., the effects of syntax and meter violations did not sum up but interacted, suggesting that the P600 effect to one or the other violation was reduced. This under-additive or reduced P600 effect from Schmidt-Kassow and Kotz (2009) fits with the current result that the P600 effect evoked by syntactic complexities was reduced in the irregular meter condition. In this light, irregular meter may have a similar impact on musicians’ syntactic integration as violated meter has on non-musicians’ syntactic integration: irregularities in the meter caused an otherwise well-formed syntactic structure to be perceived as non-preferred. This potentially implies that in spoken language, well-formedness in the metrical structure (Liberman & Prince, 1977; Prince, 1983) may play a role in syntactic comprehension in line with certain linguistic theories (Selkirk, 2011). A logical next step to pursue this possibility is to repeat the current language study with non-musicians.

### Music

We also found that the P600 effect was reduced in the irregular meter condition in music. Whereas in language this appeared to be driven by irregular meter causing the otherwise well-formed syntax to be perceived as non-preferred, in music the ERPs suggested that the non-preferred syntax was only a cost to syntactic integration when the meter was regular. Considering that meter is a component of music syntax (Lerdahl & Jackendoff, 1983), it does make sense that dispensing with part of the structure would dispense of well-formedness expectations. Upon closer inspection, this explanation is in line with previous results. In Schmuckler & Boltz (1994), though not interpreted by the authors, close inspection of the results suggests that the harmonic appropriateness of a final chord was judged as more belonging when the metrical context was not regular, e.g., with invariant timing. Similarly, Boltz (1993) observed that a regular meter was an integral component of the generation of tonal expectancies such that harmonic deviances were detected with less accuracy when presented in temporally irregular contexts. These earlier behavioral studies confirmed that tonality-syntax processing depends on the temporal organization of the musical sequence. Thus interestingly, previous ‘errors’ in judging a dissonant or tonally distant chord as appropriate in a metrically irregular context may be explained by the current results as follows: the absence of a metrically regular context may have dismantled the formation of musical syntactic preference rules, eliminating the need to syntactically integrate a non-preferred item.

It is important to note that these latter two music studies (Boltz, 1993; Schmuckler & Boltz, 1994) did not distinguish temporal context from metrical context. The stimuli in the experiments were MIDI files with exact timing, such that temporal cues were synonymous with metrical identity of phrases. Distinguishing temporal from metrical information is often not included in music research paradigms, as MIDI files with perfectly timed periodicity render the only temporal manipulations highly salient to metrical structure. This begs the question, then, whether the perception of harmonic structure is influenced by temporal or metrical characteristics, or a combination of the two. The current study was not designed to address this question, but stimuli did contain human-performed temporal fluctuations in both regular and irregular meter conditions. Thus, we avoided that metrical regularity would be synonymous with temporal periodicity. Future research on the independent effects of meter and timing can help address this knowledge gap.

Regarding the SSIRH, overall evoked responses did not confirm our hypothesis: we predicted that if syntactic integration resources are shared across domains, these resources would show a similar response to metrical regularity across domains. The current findings suggest that differences in meter-syntax alignment (Hilton & Goldwater, 2021) across domains cause different expression of the resources in the form of ERPs. However, we note that while we argue for different processing dynamics in the syntax-meter interaction here, we cannot claim that the syntactic structures were *exactly* the same across domains, merely as similar as possible in terms of more-and less syntactic complexity and more-and-less metrical regularity. In line with this, we remark that the comparability of the cross-domain cognitive processes may have been enhanced for a syntactic same-or-different discrimination task (Experiment 1) as compared to the theoretical question about relative clause attachment in language vs. tonal key assignment in music (Experiment 2). According to the SSIRH, long-term theoretical knowledge of syntactic rules is not included in the ‘shared’ cross-domain resource (Patel, 2003). Thus, cross-domain differences that we see in the ERPs may also hark back to accessing qualitatively different domain-specific episodic memory required from our task requiring theoretical knowledge of syntax. Future research would benefit from more broadly exploring the comparability of syntactic structures including expanding beyond Western language and musical cultures, as well as systematically assessing domain-specific knowledge versus cross-domain structural integration required by task demands.

### Individual differences

Individual differences were found in the music domain and significant in the accuracy of syntax discrimination. The use of factor score correlations across these studies allowed addressing a limitation in several previous musician–non-musician paradigms, namely that grouping according to musical expertise potentially masks individual differences in nuanced cognitive ability. In particular, the fact that musicians are typically associated with better temporal processing and working memory (Ho et al., 2003; Tzounopoulos & Kraus, 2009) means that improved perception of meter and syntax among musicians (e.g., Fitzroy & Sanders, 2013; Geiser et al., 2010) is indistinguishably traced back to musicianship and respectively temporal perceptual acuity or working memory. This limitation was effectively addressed here: results of Experiment 3 suggest that individual differences in perceptual processing could be attributed to cognitive resources distinctly from expertise. However, musical training may have been so influential on specific resources (like TPD) that group performance differences emerged. Thus, especially where perception of auditory signals is concerned (for tasks not requiring express knowledge such as musical theory), nuanced individual difference data is more informative than musician/non-musician grouping. In the studies mentioned above, group differences between musicians and non-musicians are potentially due to cognitive resources associated with TPD but not necessarily differences associated with VWM.

## Conclusion

Across behavioral and EEG studies, our findings generally support theories of domain-general syntactic processing. They also highlight the different roles of meter in language and music, where meter in music is more closely tied to syntax than in language. ERPs revealed that in language, irregular meter seems to disrupt the perception of syntactic well-formedness, even when the syntactic content is preferred in a regular meter context. In music, it seems that a regular meter may be a prerequisite to create stable syntactic expectations and that without it, complex harmony does not elicit any additional processing costs. These domain-differences highlight the importance of meter-syntax alignment in language and music, and suggest that future research in cross-domain syntax processing should account for this alignment in participants’ language and musical culture. Moreover, the individual differences suggest that musical ability and expertise is an important aspect to be studied further: the behavioral experiment on syntax discrimination tested musicians and non-musicians while the ERP experiment on syntactic integration necessarily tested musicians only. Thus, future studies should investigate how non-musicians exhibit language syntax preference in regular and irregular meter contexts. Individual difference findings warrant future investigations that more closely map cognitive contributions in time and pitch discrimination abilities to syntactic processing across domains, preferably in populations with a wide range of verbal- and musical abilities.

## Footnotes

1 Rather, the metrical stress placed on the wrong syllable can disrupt syntax processing (e.g., Schmidt-Kassow & Kotz, 2009).

1 We focus here on Western tonal music and stress-timed languages such as English, Dutch, and German (Abercrombie, 2022)—the language of the current study—but hope to broaden the scope to other musical cultures and language classes in the future.

1 We refer to stressed-timed languages such as English and German; metrical theory that describes multiple rhythm classes may be found in (Hayes, 1995).

1 Although there is an exception to this, for example a tendency in baroque music to shift in the final moments of a piece from a major to a minor key, called a “Picardy third” (Piston, 1987).

## Notes

### Competing Interest Statement

The authors have declared no competing interest.

